# The inhibition of automatic imitation: a meta-analysis and synthesis of fMRI studies

**DOI:** 10.1101/334938

**Authors:** Kohinoor M. Darda, Richard Ramsey

## Abstract

Humans copy other people without their conscious awareness, a behaviour known as automatic imitation. Although automatic imitation forms a key part of daily social interactions, we do not copy other people indiscriminately. Instead, we control imitative tendencies by prioritising some actions and inhibiting others. To date, neuroimaging studies investigating the control of automatic imitation have produced inconsistent findings. Some studies suggest that imitation control relies on a domain-specific neural circuit related to social cognition (the theory-of-mind network). In contrast, other studies show engagement of a domain-general neural circuit that is engaged during a diverse range of cognitive control tasks (the multiple demand network). Given the inconsistency of prior findings, in the current paper we avoided problems associated with interpreting individual studies by performing a meta-analysis. To do so, we used a multi-level kernel density analysis to quantitatively identify consistent patterns of activation across functional magnetic resonance imaging studies investigating the control of imitation. Our results show clear and consistent evidence across studies that the control of automatic imitation is guided by brain regions in the multiple demand network including dorsolateral frontoparietal cortex. In contrast, there was only limited evidence that regions in the theory of mind network were engaged. Indeed, medial prefrontal cortex showed no consistent engagement and right temporoparietal junction engagement may reflect spatial rather than imitative control. As such, the current meta-analysis reinforces the role of domain-general control mechanisms and provides limited evidence in support of the role of domain-specific processes in regulating imitative tendencies. Consequently, neurocognitive models of imitation need updating to place more emphasis on domain-general control mechanisms, as well as to consider more complex organisational structures of control, which may involve contributions from multiple cognitive systems.

## Introduction

The involuntary tendency of human beings to imitate others’ gestures, speech patterns, and postures, is known as automatic imitation (Heyes, 2011). It has been suggested that such automatic imitative behaviour functions as a “social glue” as it increases pro-social behaviour, positive rapport, feelings of affiliation and liking between interacting partners (Kavanagh & Winkielman, 2016; van Baaren, Janssen, Chartrand, & Dijksterhuis, 2009; Lakin & Chartrand, 2003; Chartrand & Bargh, 2009; van Baaren, Holland, Steenaert, Van Kippenberg, 2003). Given the influence of imitation on strengthening social bonds, researchers have started to investigate the psychological and biological mechanisms that underpin imitation. For example, over the last 20 years, researchers have used functional magnetic resonance imaging (fMRI) in order to better understand the neural underpinnings of the control of automatic imitative tendencies. However, these studies have provided mixed findings regarding the contributions of domain-general or domain-specific neural networks in imitation control. The current paper, therefore, meta-analyses fMRI studies to date on the control of automatic imitation in order to provide a combined quantitative estimate of the extant evidence of many individual studies (Lipsey & Wilson, 2001).

In the last two decades, automatic imitation has been widely studied with an attempt to interconnect different disciplines like cognitive science, social psychology, evolutionary biology, and cognitive neuroscience (Prinz & Meltzoff, 2002; Chartrand & Bargh, 1999; Byrne & Russon, 1998). This convergence across multiple disciplines has allowed for a range of perspectives on imitation to emerge in which theory and empirical data can strengthen each other. In social psychology, automatic imitation has been studied in naturalistic social interactions (Chartrand & Lakin, 2013). Along with functioning as a “social glue,” research performed in more naturalist settings suggests that imitation behaviour is also moderated by other variables including, but not limited to, personality variables, self-construal, goal to affiliate or disaffiliate, cultural and social contexts, as well as the similarity, familiarity, and status of the person being imitated (Chartrand & Lakin, 2013; Caspers et al., 2010; Duffy & Chartrand, 2015).

Even though automatic imitation seems to be an important behaviour that facilitates social interactions, we do not always copy others’ behaviours. In many situations, imitation can be maladaptive, and it is essential to circumvent the tendency to automatically imitate (Cross & Iacoboni, 2014; Cross, Torrisi, Losin, & Iacoboni, 2013; van Schie, van Waterschoot, & Bekkering, 2008; Newman-Norlund, van Schie, van Zuijlen, & Bekkering, 2007). This need to regulate imitative tendencies indicates the existence of a selection mechanism that inhibits unwanted actions, and prioritises alternatives (Brass et al., 2009). Thus, imitation control can be divided into at least two component processes – action representation and action selection. We observe an interaction partner and their actions (representation), and then select the action that needs to be executed (selection).

In contrast to social psychology approaches, researchers in the field of cognitive psychology and neuroscience have generally used computer-based reaction-time (RT) measures of the inhibition of automatic imitation (Brass et al., 2000; Stürmer et al., 2000). One of the most commonly used tasks in this field consists in making finger movements while simultaneously observing a compatible or incompatible finger movement (Brass et al., 2000). For example, participants may be asked to make a finger movement in response to an imperative cue i.e. they are instructed to lift their index finger when they see a number ‘1’ on screen, and their middle finger when they see a number ‘2.’ Simultaneously, participants also observe a task-irrelevant index or middle finger movement, which is compatible or incompatible with their own response. Other variants of this task include hand opening and closing movements instead of finger movements (Press et al., 2005; Wang et al., 2011) or pre-specifying the participant’s response before the imperative cue (i.e. participants are asked to always lift their index finger when they see a finger movement; Brass, Bekkering & Prinz, 2001; Heyes et al., 2005). In these variants as well, participants observe a hand or finger movement which is compatible or incompatible with their own response. Irrespective of the task used, greater cognitive resources are required when inhibiting movements incompatible to one’s own responses, thus leading to greater RTs (Heyes, 2011; Brass & Heyes, 2005). The difference between the incompatible and compatible conditions (referred to as the general compatibility effect) is said to be a measure of imitation control (Heyes et al., 2005; Heyes, 2011).

To date, a number of neuroimaging studies have investigated the neural mechanisms of imitation control using RT paradigms. However, the evidence demonstrating the extent to which RT paradigms of imitation control engage domain-general or domain-specific neural networks is mixed. Domain-specific processes operate on particular types of stimuli or aspects of cognition, while domain-general processes operate across a range of stimuli and tasks (Barett, 2012; Spunt & Adolphs, 2017). One of the prevailing theories of automatic imitation proposes that imitation control relies on a domain-specific neural circuit related to social cognition (Brass et al., 2009). This “specialist” theory has gained traction with evidence from patient and neuroimaging data pointing to the engagement of two key candidate regions – the anterior medial prefrontal cortex (mPFC) and the right temporoparietal junction (rTPJ) (Brass & Heyes, 2005; Brass et al., 2009). For example, mPFC and rTPJ have been engaged in human brain imaging investigations of imitation inhibition (Brass et al., 2001; 2005; 2009; Spengler et al., 2009; Wang et al., 2011). Brass and colleagues further proposed a dissociation of roles for the mPFC and rTPJ during imitation control - the rTPJ distinguishes between self- and other-generated actions, and the mPFC enforces the self-generated action when faced with conflict from an action representation generated by another agent (Brass et al., 2009). In addition, patients with frontal lobe lesions show disrupted imitation inhibition behaviour (Brass et al., 2003; Spengler et al., 2010) and an increased tendency to automatically imitate even when they are clearly instructed to not do so (Lhermitte et al., 1986). More evidence for the involvement of rTPJ comes from neuro-stimulation studies: inhibiting the activity in the rTPJ by transcranial magnetic stimulation (TMS) interfered with imitative responses impairing imitation inhibition (Hogeveen et al., 2014; Sowden & Catmur, 2015). Irrespective of the method used, it is worth noting that, to date, there have only been a small number of studies implicating mPFC and rTPJ in the control of imitation. Moreover, these studies have used relatively small sample sizes between 10 and 25 participants and there have been few, if any, direct replications. Therefore, the sum total of evidence for mPFC and rTPJ engagement during imitation control is suggestive rather than compelling.

Along with imitative control, neuroimaging findings suggest mPFC and rTPJ are also engaged in a variety of socio-cognitive tasks that are associated with theory of mind, including distinguishing between self from other, perspective taking, as well as attributing beliefs, desires and attitudes to others (ToM; Gallagher et al., 2000; Amodio & Frith, 2006; Ruby & Decety, 2001; Aichhorn et al., 2006; Decety et al., 2002; Santiesteban et al., 2012; Brass et al., 2009; Spengler et al., 2010). Based on these findings, self-other control processes have thus been proposed as a candidate mechanism for a range of socio-cognitive functions. For example, it is important to inhibit one’s own perspective or mental state and enhance that of the other when empathising with others, taking their perspective, or engaging a successful theory-of-mind (de Guzman et al., 2016; Sowden & Shah, 2014). Further, atypical self-other control has been linked to disorders characterised by social dysfunction including autism and schizophrenia (Cook and Bird, 2012; Ferri et al., 2012). Overall, this evidence suggests that in imitation control, it is crucial to inhibit the representation of the other’s action, and enforce your own, and this mechanism is guided by a domain-specific neural circuit unique to social cognition (Brass et al., 2009).

In contrast to this “specialist” view of imitation control, however, “generalist” theories of imitation suggest that the inhibition of automatic imitation does not differ from any other pre-potent tendencies or general cognitive functions (Heyes, 2011; Cooper et al., 2013). Multiple cognitive control tasks like the Flanker, Stroop, and Simon tasks, which require the inhibition of automatic overlearned response tendencies, have been found to engage a domain-general control network identified in the dorsolateral fronto-parietal cortices (Aron et al., 2014; Bunge et al., 2002; Hazeltine et al., 2007; Nee et al., 2007; Wager et al., 2005). This network is also called the multiple demand (MD) network as it is engaged across a diversity of mental operations (Duncan et al., 2010). Across studies that investigate imitation inhibition, some have found engagement of the mPFC and rTPJ (Brass et al., 2001; 2005; 2009; Spengler et al., 2009), whereas others show engagement of the MD network (Bien, Roebroeck, Goebel, & Sack, 2009; Crescentini, Mengotti, Grecucci, & Rumiati, 2011; Cross & Iacoboni, 2013; Mengotti, Corradi-Dell’Acqua, & Rumiati, 2012; Marsh et al., 2016). However, most previous fMRI studies have been limited by low statistical power and small sample sizes. More recently, a multi-experiment study using larger sample sizes (N=28, N=50) and a functional region of interest (fROI) approach that bolsters statistical power and functional sensitivity has shown that imitation control engages only the MD network, and not mPFC or rTPJ (Darda, Butler & Ramsey, 2018). Indeed, even with an *a priori* power analysis ensuring 80% power to detect medium effect sizes, Darda and colleagues (2018) did not even find a directional trend to suggest that the ToM network was directly engaged during imitation control.

As mentioned before, imitation control can be divided into at least two component processes – action representation and action selection. The above review of literature suggests two possible neural mechanisms as being key to action selection during imitation control. On one hand, during imitation control, the neural representation generated by the observed person’s action is inhibited, and the self-generated action is selected and enforced and this selection mechanism engages a domain-specific neural network i.e. the mPFC and rTPJ. On the other hand, the selection mechanism may be guided by a domain-general neural network i.e. the MD network. In both possible mechanisms, the input is the same i.e. the observed person and action may engage domain-specific socio-perceptual neural circuits. However, the difference lies in the selection or control mechanism that underlies the inhibition of automatic imitative tendencies which finally leads to consequent behaviour (see graphical representation in Figure 1).

**Figure 1.**
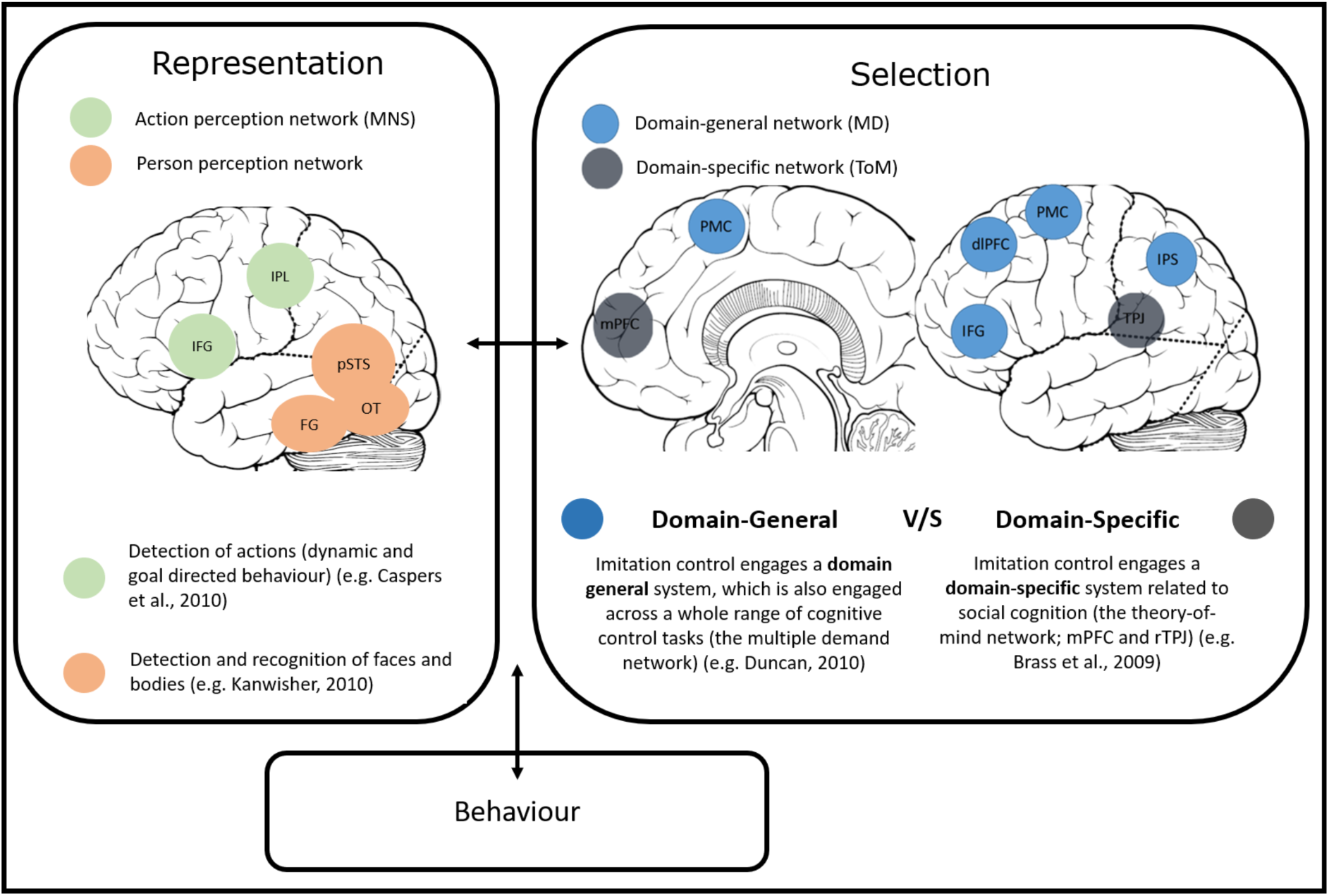
Brain networks associated with the control of automatic imitation. This graphical representation divides imitation control into two constituent processes – representation of the person and their action, and the selection (control) of the right action to be executed. In the context of automatic imitation, the representation system consists in face, body, biological motion, and action perception. The neural substrates for person and action perception span the fusiform gyrus, occipitotemporal cortex, and posterior superior temporal sulcus, as well as the mirror neuron system (Kanwisher, 2010; Caspers, et al., 2010). The control or selection system consists in a brain network that is either domain-general (i.e. the multiple demand network) or domain-specific (i.e. the theory-of-mind network). N.B. Abbreviations: MNS = mirror neuron system; IPL = inferior parietal lobule, IFG = inferior frontal gyrus; pSTS = posterior superior temporal sulcus; OT = occipito-temporal cortex; FG = fusiform gyrus, MD = multiple demand network; ToM = theory-of-mind network; mPFC = medial prefrontal cortex; PMC = primary motor cortex; dlPFC = dorsolateral prefrontal cortex; TPJ = temporo-parietal junction. The bidirectional arrow “**↔**” indicates links between the different nodes of imitation control.

The question of interest for the current meta-analysis, therefore, lies at the selection stage of imitation control with the evidence to date for engagement of domain-specific and domain-general neural networks being inconsistent. Even though the most statistically powerful fMRI study to date only shows the engagement of the MD network (Darda et al., 2018), the interpretation of individual studies remains limited in scope for several reasons. First, many single studies are likely to be underpowered leading to missed or spurious results (Button et al., 2013). Second, empirical work involves design choices that strongly influence results, making it harder to generalise effects across analysis pipelines and differing experimental procedures (Carp, 2012). Given the inconsistency of prior findings and the absence of a quantitative synthesis of evidence, taking a meta-analytical approach to further investigate the neural basis of imitation has many benefits (Cumming, 2014). As such, by means of a meta-analysis, the current paper enables the detection of consistent patterns of activation across studies.

In order to quantify the consistency and specificity of regional activation for imitation control across studies, we performed a multi-level kernel density analysis (MKDA; see Methods and Materials for details). We included all fMRI studies (N=12) investigating imitation control using the RT measure of imitation inhibition (see Table 1). Our primary measure aimed to quantify the consistency of region engagement across studies with particular focus on the engagement of the ToM network and the MD network. The dependent variable was the blood oxygen level dependent (BOLD) response measured in the included fMRI studies. Given the prior mixed findings across studies, this meta-analysis aimed to quantify the extent to which ToM, MD or both neural networks may be engaged when during the inhibition of automatic imitation.

**Table 1.**
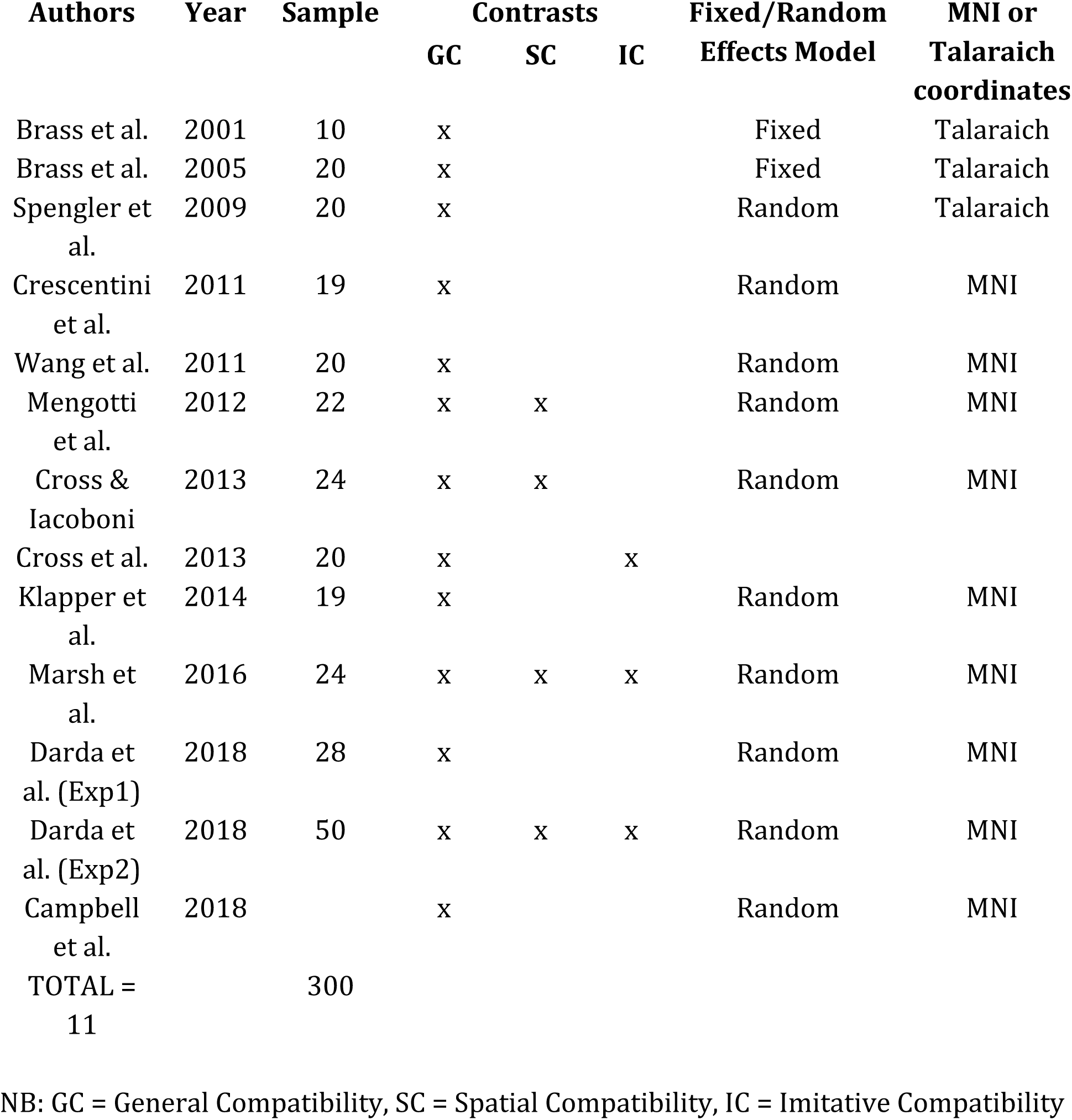
Data extracted from the studies included in the meta-analysis.

We also ran two more exploratory analyses, which were based on a small subset of the total studies. The most common measure of imitation inhibition, the general compatibility effect, also includes a spatial component (Heyes, 2011). In order to measure imitative compatibility more specifically, therefore, imitative and spatial effects need to be dissociated (Gowen et al., 2016; Boyer et al., 2012; Catmur & Heyes, 2011). However, only a few fMRI studies have measured the imitative compatibility effect independent of the spatial component (Darda et al., 2018; Marsh et al., 2016; Cross et al., 2013). This makes it difficult to interpret the roles of the ToM (mPFC and rTPJ) and MD networks in imitation control – their engagement could reflect both social (imitative) and/or non-social (spatial) control. Indeed, the rTPJ has been previously associated with orienting to both social and non-social stimuli (Corbetta et al., 2008; Thiel et al., 2004). Thus, given that only a few studies have dissociated between imitative (N=3) and spatial compatibility (N=4) effects, we also ran two further exploratory MKDAs in order to quantify consistency of patterns across studies for both imitative and spatial compatibility effects. Indeed, given the low number of studies included in the secondary analyses, these results provide only suggestive, and not compelling, evidence regarding the role of the MD and ToM networks in imitative and spatial control.

## Methods and Materials

### Literature search and data collection

In the current paper, we follow recent guidelines put forward for meta-analysing neuroimaging studies (Muller et al., 2018). FMRI studies exploring the inhibition of automatic imitative tendencies were searched for on the online database PubMed, as well as the article search engine Google Scholar. Combinations of keywords including ‘imitation inhibition,’ ‘fMRI,’ ‘imitation,’ ‘automatic imitation,’ and ‘imitation control’ were used to identify relevant literature (prior to January 2019). A total of 15 studies were found. We rejected studies if the primary method of investigation was not fMRI, if the study did not report results in stereotactic coordinate space (either Montreal neurological Institute (MNI) or Talaraich coordinates) (N=1; Bien et al., 2009), if reported results were based on region-of-interest (ROI) analyses, and the study did not report whole-brain analysis coordinates either in the main article or in supplementary materials (or we could not obtain them from the authors) (N=1; Brass et al., 2009), and if the study involved children or atypical populations (and the coordinates for controls were not reported separately) (N=1; Spengler, Bird, & Brass, 2010).

A wide variety of contrasts are used in studies that investigate the inhibition of automatic imitation. However, in order to minimise heterogeneity, studies that used a paradigm that was not based on or was not conceptually similar to the Brass et al. (2000) paradigm for measuring inhibition of automatic imitation were also excluded. Thus, 12 studies with a total of 300 participants were included in the meta-analysis (see Table 1).

Even though our main analysis was on the general compatibility effect, we also ran two separate meta-analyses for spatial and imitative compatibility. Table 2 summarises the contrasts used in the current meta-analysis for general, spatial, and imitative compatibility effects. A total of 13 contrasts across 12 studies with 142 foci were used for general compatibility, 4 contrasts across 4 studies with 42 foci were used for spatial compatibility, and a total of 3 contrasts across 3 studies with 20 foci were used for imitative compatibility.

**Table 2.** Contrasts used in the meta-analysis for general, spatial, and imitative compatibility.

| **Authors** | **Year** | **General Compatibility** | **Spatial Compatibility** | **Imitative Compatibility** |
| --- | --- | --- | --- | --- |
| Brass et al. | 2001 | General Incompatible > General Compatible |  |  |
| Brass et al. | 2005 | General Incompatible > General Compatible |  |  |
| Spengler et al. | 2009 | General Incompatible > General Compatible |  |  |
| Crescentini et al. | 2011 | General Incompatible > General Compatible |  |  |
| Wang et al. | 2011 | General Incompatible > General Compatible* |  |  |
| Mengotti et al. | 2012 | Non-specular > Specular^ | Spatially Incompatible > Spatially Compatible |  |
| Cross & Iacoboni | 2013 | General Incompatible > General Compatible | Spatially Incompatible > Spatially Compatible |  |
| Cross et al. | 2013 | General Incompatible > General Compatible |  | General Compatibility > Spatial Compatibility |
| Klapper et al. | 2014 | General Incompatible > General Compatible* |  |  |
| Marsh et al. | 2016 | General Incompatible > General Compatible | Spatial Incompatible > Spatially Compatible | Imitatively Incompatible > Imitatively Compatible |
| Darda et al. (Exp1) | 2018 | General Incompatible > General Compatible |  |  |
| Darda et al. (Exp2) | 2018 | General Incompatible > General Compatible | Spatial Incompatible > Spatial Compatible | Imitative Incompatible > Imitatively Compatible |
| Campbell et al. | 2018 | General Incompatible > General Compatible |  |  |
| **No. of studies** |  | 12 | 4 | 3 |
| **No. of contrasts** |  | 13 | 4 | 3 |
| **No. of foci** |  | 142 | 42 | 20 |

Table 2 shows the contrasts used in the current meta-analysis, and the number of contrasts, foci, and studies for each compatibility type (general, spatial, and imitative).
\*Collapsed across conditions; for Wang et al., 2011: collapsed across direct and averted gaze, for Klapper et al., 2014: collapsed across belief (motion-capture, computer animation) and form (human, non-human).
^Non-specular > Specular i.e. {(spatially incompatible and imitatively compatible) + (imitatively incompatible and spatially compatible) > general compatible}}

### Data analysis

All analyses in the current paper were performed in MatlabR2015b (Mathworks, Naticks, MA) using the MKDA toolbox developed by Wager et al., 2007; http://wagerlab.colorado.edu). MKDA is an analysis technique that uses a random effects model to assess convergence across studies. This allows for assessing convergence across studies as opposed to between individual foci (as implemented in classical meta-analysis techniques that use fixed effects analyses). Thus, results are not biased by a small number of individual studies. Further, each contrast is weighted by the sample size and study quality (i.e. whether the study used fixed or random effects model; Wager et al., 2007; Kober and Wager, 2010).

MKDA was performed on all three compatibility types separately. Before performing the analyses, we extracted the following information from each study and included it in our database: authors, year of publication, sample size, task contrasts, fixed or random effects model, and MNI or Talaraich co-ordinates. Co-ordinates reported in Talairach space were converted to MNI stereotactic space using Lancaster transformation (tal2icbm transform; Lancaster et al., 2007). Peak coordinates from each contrast map were then convolved with a 10mm spherical kernel in order to create a contrast indicator map (CIM). The resulting voxels within 10 mm of the peak were deemed “significant” and given a value of one; other voxels were given a value of zero which indicated no significant effect. A density map was then created by averaging the indicator maps, weighted by sample size, and whether the study used a fixed or random effects model. More specifically, as recommended by Wager and colleagues (Wager, Lindquist, & Kaplan, 2007), this density map was weighted by the square root of the sample size of the study, and then multiplied by an adjustment factor of 1 for random effects analysis, and .75 for a fixed effects analysis.

Each voxel of the density map was given a density statistic P. P stands for the proportion of contrasts included in the analysis that show activity within 10mm of the voxel. A Monte Carlo simulation (with 5000 iterations) was then carried out in order to identify voxels that had a P-statistic that was higher than the frequency predicted by chance. This was tested against the null hypothesis that activated regions in the resulting pairwise contrast maps (from the 5000 iterations) were randomly distributed across the brain. To test for the significance of the cluster size, a similar procedure was used. This allowed for the identification of a threshold for cluster size at which a specific number of voxels needed to be activated contiguously so that the cluster could be deemed significant.

In order to maximise sensitivity in testing our hypotheses, we report results using two thresholding techniques. One thresholding technique is based on height and the other is based on cluster size. For the weighted P-statistic (height-based threshold), the family wise error (FWE) corrected threshold is the proportion of studies which yielded activity within 10 mm of a voxel that showed a higher P-statistic than the maximum P-statistic across 95% of the Monte Carlo maps. For the cluster size threshold, the FWE corrected threshold is the contiguous voxels observed at two different thresholds (p<.001 and p<.01) whose cluster size is more than the extent of clusters found across 95% of the Monte Carlo maps. We use two cluster-based thresholds in order to also detect regions that show a lower response in magnitude over a larger cluster size both at more stringent (p<.001) and less stringent (p<.01) thresholds. Voxels that exceed the height-based threshold in our analysis appear on the resulting maps in Figure 2 in yellow, and those that exceed the cluster extent-based threshold appear in orange (p<.001) and red (p<.01).

**Figure 2.**
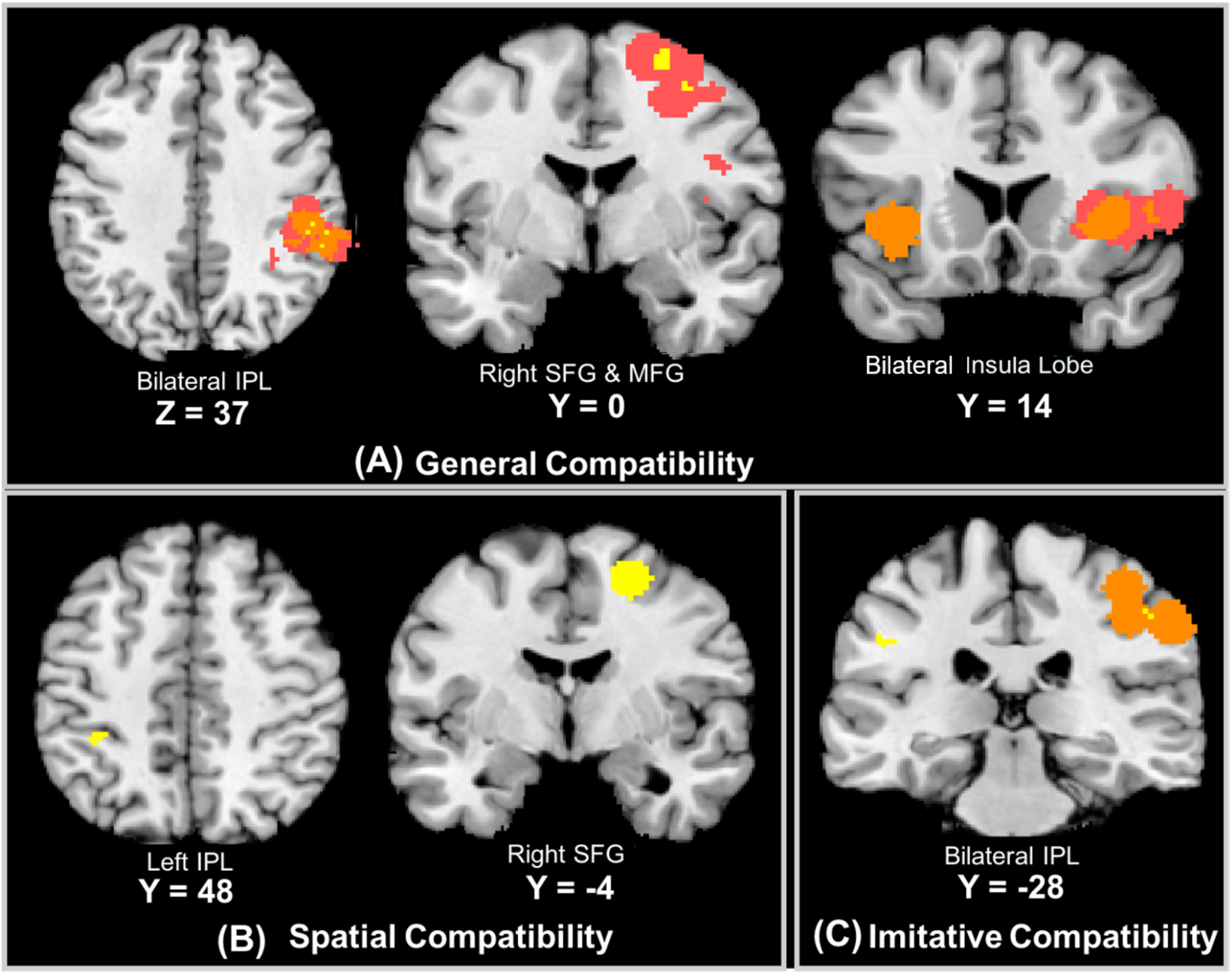
Consistency of brain activation from the MKDA Analyses. Brain areas that are consistently engaged for general compatibility (A), spatial compatibility (B), and imitative compatibility (C). Voxels that exceed the height-based threshold (p<.05, FDR corrected) in our analysis appear in yellow, and those that exceed the cluster extent-based threshold appear in orange (p<.001) and red (p<.01).

In the resulting table (Table 3), peak activation foci that pass the height-based threshold are reported. If activations do not pass the height-based threshold, foci of the cluster-extent-based thresholding are reported. The number of voxels in each cluster that survived height-based and/or extent-based thresholding is also reported. Resulting coordinates were localised using the SPM Anatomy Toolbox (Eickhoff et al., 2005). The database of co-ordinates, and code used to perform the meta-analysis are available online (https://osf.io/dbuwr/).

**Table 3.** Areas consistently activated for general compatibility, spatial compatibility, and imitative compatibility.

| GENERAL COMPATIBILITY |  |  |  |  |  |  |
| --- | --- | --- | --- | --- | --- | --- |
| Region | MNI |  |  | Maximum P | No. of Voxels | Threshold |
|  | x | y | z |  |  |  |
| **Right TPJ** | 60 | -46 | 22 | 0.42 | 1 | ^** |
| **Right TPJ** | 56 | -46 | 32 | 0.41 | 1 | ^** |
| **Right supramarginal gyrus** | 56 | -36 | 36 | 0.44 | 9 | ^** |
| **Right IPL** | 60 | -34 | 34 | 0.43 | 2 | ^** |
| **Right IPL** | 52 | -30 | 38 | 0.37 | 5 | ^** |
| **Right MFG** | 34 | 0 | 54 | 0.45 | 15 | ^* |
| **Right SFG** | 26 | -2 | 64 | 0.37 | 46 | ^* |
| **Left Insula Lobe** | -36 | 14 | 0 | 0.32 | 453 | ** |
|  | -34 | 12 | -2 |  | 213 |  |
|  | -36 | 18 | 2 |  | 240 |  |
| **Right Insula Lobe** | 38 | 16 | 4 | 0.36 | 405 | ** |
|  | 34 | 18 | 0 |  | 172 |  |
|  | 46 | 12 | 2 |  | 75 |  |
|  | 38 | 16 | 6 |  | 158 |  |
| **Right IFG** | 46 | 14 | 10 | 0.28 | 1269 | * |
|  | 44 | 16 | -4 |  | 74 |  |
|  | 28 | 24 | -4 |  | 44 |  |
|  | 32 | 12 | 0 |  | 41 |  |
|  | 46 | 22 | 2 |  | 113 |  |
|  | 56 | 10 | 2 |  | 116 |  |
|  | 28 | 20 | 6 |  | 105 |  |
|  | 36 | 26 | 6 |  | 66 |  |
|  | 56 | 16 | 6 |  | 128 |  |
|  | 52 | 12 | 14 |  | 219 |  |
|  | 48 | 2 | 22 |  | 92 |  |
|  | 40 | 8 | 22 |  | 132 |  |
|  | 50 | 8 | 28 |  | 139 |  |
| **SPATIAL COMPATIBILITY** |  |  |  |  |  |  |
| **Left IPL** | -36 | -40 | 48 | 0.78 | 8 | ^ |
| **Right superior frontal gyrus** | 24 | -4 | 58 | 0.78 | 190 | ^ |
|  | 24 | -6 | 54 |  | 56 | ^ |
|  | 24 | -4 | 60 |  | 134 | ^ |
| **IMITATIVE COMPATIBILITY** |  |  |  |  |  |  |
| **Left supramarginal gyrus/ IPL** | -48 | -28 | 34 | 0.73 | 11 | ^ |
| **Right supramarginal gyrus/ IPL** | 48 | -26 | 44 | 0.73 | 18 | *^ |
|  | 52 | -30 | 42 |  | 7 |  |
|  | 46 | -26 | 46 |  | 11 |  |

Table 3 shows areas consistently activated for general compatibility, spatial compatibility, and imitative compatibility. Maximum P stands for the maximum proportion of studies exhibiting the effect at the peak density weighted by sample size. MNI = Montreal Neurological Institute (MNI) standard stereotaxic space coordinates. The voxel size is $2 \times 2 \times 2 \text{ mm}^3$ .
\*Clusters withstanding $p < .01$ cluster extent-based threshold.
\*\*Clusters withstanding $p < .001$ cluster extent-based threshold.
^Clusters withstanding the height-based threshold.

## Results

For the general compatibility effect, across 13 contrasts from 12 studies, consistent activation was found in right inferior parietal lobule, right supramarginal gyrus, right superior temporal gyrus, and right temporo-parietal junction (see Table 3; Figure 2A). These clusters survived both height-based and the more stringent extent-based thresholding (p<.001). Activation was also found in right superior frontal gyrus, and right middle frontal gyrus, which survived both height and the less stringent extent-based thresholding (p<.01). Activation in the left and right insula survived the more stringent extent-based threshold (p<.001), but not the height-based threshold. Activation in the right IFG survived the less stringent extent-based threshold (p<.01) but not the height-based threshold.

We ran two further MKDAs separately for spatial and imitative compatibility. For spatial compatibility, across 4 contrasts from 4 studies, we found consistent activation that withstood the height-based thresholding in the left IPL and the right SFG (see Table 3; Figure 2B). No regions withstood cluster-based thresholding. For imitative compatibility, across 3 contrasts from 3 studies, we found consistent activation in the left IPL that survived height-based (see Table 3; Figure 2C). Activation was also found in the right IPL, which withstood height-based as well as the less stringent extent-based thresholding (p<.01).

These density maps showing regions that withstood both height and/or cluster-extent thresholding for each compatibility type were then overlaid with the ToM and MD network masks separately. The ToM network mask consisted of four parcels including the dorsal, medial, and ventral medial prefrontal cortex (DMPFC, MMPFC, VMPFC), and the right temporo-parietal junction (rTPJ), which have previously been implicated in mentalising or theory-of-mind (Dufour et al., 2013). For the MD network mask, 16 parcels were used which included areas in bilateral superior and inferior parietal lobules (SPL, IPL), intraparietal sulcus (IPS), inferior and middle frontal gyrus (IFG, MFG), precentral gyrus (PrecG), insula (Ins), and the supplementary motor area (SMA) (available at: https://evlab.mit.edu/funcloc/download-parcels). Overlay of the density maps with the ToM and MD network masks allowed for identification of overlap between regions that were consistently activated in the MKDA and the ToM and MD networks (Figure 3). For all compatibility types (general, imitative and spatial), all regions that passed height or extent-based thresholding overlapped with regions in the MD network (Figure 3A). Additionally, one cluster, which showed consistent activation for general compatibility, also overlapped with the right TPJ in the ToM network (Figure 3B). There was no overlap with the mPFC node of the ToM network for any compatibility type.

**Figure 3.**
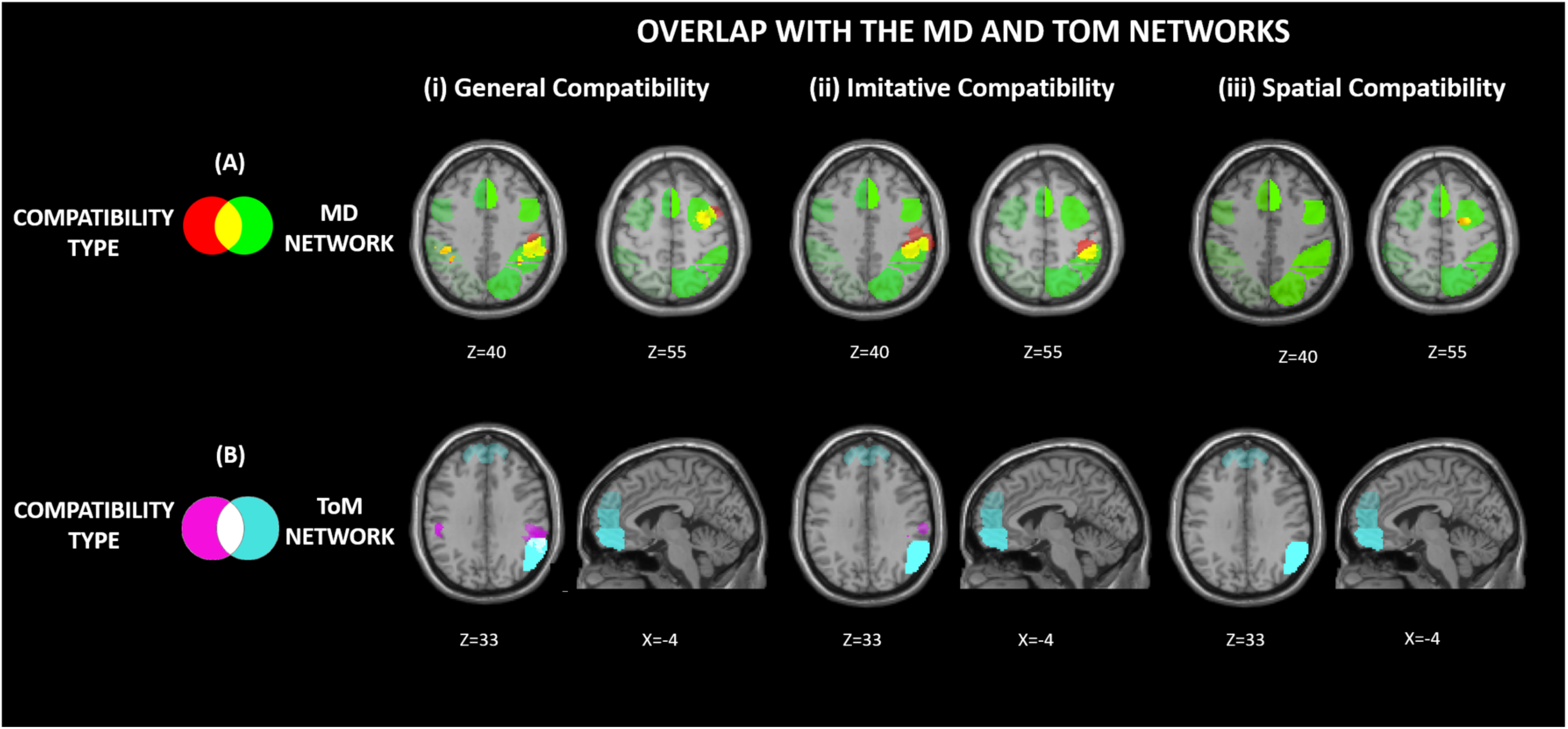
Overlay of the MKDA maps with the ToM and MD network masks. Overlay of the density maps with the ToM and MD network masks allowed for identification of overlap between regions that were consistently activated in the MKDA and the ToM and MD networks. For all compatibility types (general, imitative and spatial), all regions that passed height or extent-based thresholding overlapped with regions in the MD network (A). Additionally, one cluster, which showed consistent activation for general compatibility, also overlapped with the right TPJ in the ToM network (B). There was no overlap with the mPFC node of the ToM network for any compatibility type.

In order to break down the role of the right TPJ in general compatibility, we performed a further, more exploratory analysis. We compared peak coordinates from prior studies with a right TPJ mask, which has been previously implicated in theory-of-mind (Dufour et al., 2013). To do so, the ToM network mask for rTPJ was overlaid with the contrast indicator maps of all studies used for imitative (N=3) and spatial (N=4) compatibility. The contrast indicator maps include 10mm spherical kernels around peak coordinates of each contrast. This allows coordinates from prior imitative and spatial compatibility contrasts to be displayed without any thresholding restrictions and overlaid with the rTPJ node of the ToM network. Figure 4 shows overlap between contrast indicator maps for general compatibility and spatial compatibility with the right TPJ node of the ToM network mask. By contrast, there is no overlap between contrast indicator maps for imitative compatibility and the same right TPJ mask.

**Figure 4.**
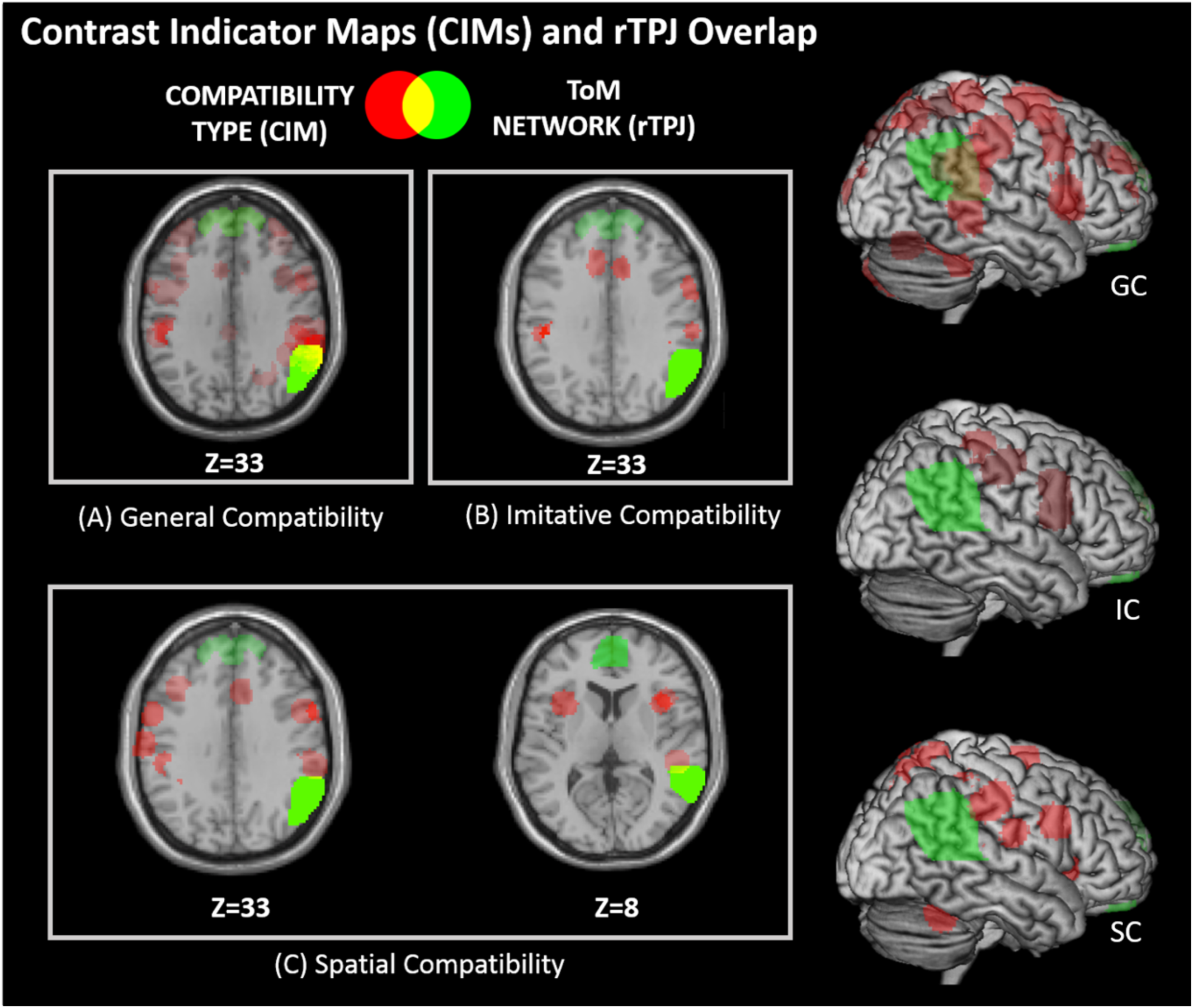
Overlay of Contrast Indicator Maps with rTPJ. The ToM network mask for rTPJ overlaid with the contrast indicator maps of all studies used for general (N=12; A) imitative (N=3; B) and spatial (N=4; C) compatibility. There was overlap between contrast indicator maps for general compatibility and spatial compatibility with right TPJ. There is no overlap between contrast indicator maps for imitative compatibility and the same right TPJ mask. Abbreviations: IC = Imitative Compatibility; SC = Spatial Compatibility, GC = General Compatibility

## Discussion

In the current paper, we performed a meta-analysis of fMRI studies in order to quantify the consistency and specificity of regional activation during the inhibition of automatic imitation. Our results supported a “generalist” view of imitation control – we found clear engagement of dorsolateral frontoparietal cortices when observing an action that conflicted with a current motor intention. These regions overlapped with regions associated with the MD network. We found less evidence for a “specialist” view of imitation control, which relies on the ToM network. Indeed, there was no engagement of mPFC across studies and there was no clear evidence regarding the engagement of rTPJ; there was only suggestive evidence that it may reflect spatial rather than social control. Thus, our results provide unambiguous support for the engagement of a domain-general neural network during the control of imitation, and only limited evidence for the engagement of a domain-specific neural network that is tied to social cognition.

Studies investigating the neural correlates of imitation control have to date shown mixed evidence for the engagement of domain-general and domain-specific neural networks in imitation inhibition. While some studies have found engagement of the mPFC and rTPJ (Brass et al., 2001; 2005; 2009; Spengler et al., 2009), others show engagement of the MD network (Bien, Roebroeck, Goebel, & Sack, 2009; Marsh et al., 2016; Darda et al., 2018). The current MKDA demonstrated that brain regions in the multiple demand network are reliably and consistently engaged across studies that investigate imitation inhibition using the general compatibility effect. Brain regions in the MD network are also engaged for imitative (bilateral IPL) and spatial compatibility effects (left IPL, right SFG). Thus, our findings suggest that brain regions that are engaged across a range of cognitive control tasks are also reliably engaged when controlling the automatic tendency to imitate others, as measured by general and imitative compatibility effects.

Evidence supporting the engagement of a domain-specific neural circuit that is central to social cognition and includes mPFC and rTPJ was less consistent in the current meta-analysis. Brass et al. (2009) proposed that the rTPJ was involved in distinguishing between self- and other-generated actions, whereas the mPFC was engaged when enforcing the correct action. However, the current MKDA did not find any evidence of anterior mPFC engagement for either general, spatial, or imitative compatibility effects. An absence of mPFC engagement for imitation control across studies is thus inconsistent with the hypothesis that a specific neural system related to social cognition is also engaged during the inhibition of automatic imitation (Brass et al., 2009).

In contrast to the results reported in mPFC, across studies investigating imitation inhibition as measured by the general compatibility effect, the current meta-analysis found engagement of rTPJ. However, it is difficult to interpret the role of rTPJ in imitation control for at least two reasons. First, the general compatibility effect is a product of both spatial and imitative effects, which makes it hard to interpret in a straightforward manner. Second, rTPJ is involved in both social and non-social processes, which makes it a functionally heterogenous region (Corbetta et al., 2008; Krall et al., 2015, 2016; Lee & McCarthy, 2016; Schuwerk et al., 2017).

Further, in the current meta-analysis, across 12 studies, we find engagement of rTPJ for the general compatibility effect. However, it is important to distinguish between a synthesis of evidence based on a descriptive approach, and a quantitative meta-analysis (Gigerenzer, 2018). To date, 14 fMRI studies have investigated imitation control by measuring the general compatibility effect (we excluded Bien et al., 2009 and Brass et al., 2009 in the meta-analysis, see Methods). Out of the 14 studies, only 4 studies report the engagement of rTPJ for the general compatibility effect (see Darda et al., 2018, Table 1 for more details). Thus, we find that only 28.6% of fMRI studies on imitation control to date (4/14) show any evidence in support of a role of rTPJ in imitation control, and these studies do not dissociate between spatial and imitative effects.

In the largest and most sensitive fMRI study of imitation inhibition to date, Darda and colleagues (2018) showed no engagement of rTPJ for the imitative compatibility effect, but engagement of rTPJ for general and spatial compatibility effects. Similarly, in the current meta-analysis, when we explored the unthresholded spatial and imitative compatibility effect maps separately, there was partial overlap between the spatial compatibility effect and rTPJ, but no overlap between the imitative compatibility effect and rTPJ. Given that the small number of fMRI studies investigating spatial and imitative compatibility effects separately (N=4 and N=3, respectively), the current findings need to be interpreted with caution. However, when taken together with prior findings, the results provide consistently limited evidence for the univariate engagement of rTPJ in the control of imitative tendencies. In contrast, current fMRI findings provide more evidence that rTPJ is involved in resolving spatial conflict, which is in keeping with patient work (Vallar & Perani, 1987; Valler et al., 1993), as well as evidence using spatial cueing tasks like the Posner paradigm (Posner & Cohen, 1984; Thiel et al., 2004; Corbetta et al., 2008). More recent work also suggests that rTPJ may play a more domain-general role in the process of contextual updating, acting on changing expectations after unexpected events (Geng & Vossel, 2013; Mengotti et al., 2017). Assuming that on incompatible trials expectations are violated, rTPJ may play a more generalised role of context updating in imitation and spatial control. However, irrespective of whether it plays a domain-specific or domain-general role, in the current meta-analysis, we find limited evidence for the univariate engagement of rTPJ in the control of automatic imitative tendencies.

### Limitations and alternative interpretations

Before moving on to the wider theoretical implications of these results, we first acknowledge possible limitations to the current meta-analytical approach. The current meta-analysis did not include work by Brass and colleagues (2009), which implicated rTPJ and mPFC in imitation control, due to the whole-brain data being unavailable. Nonetheless, as mentioned before, only 28.6% (4/14) of fMRI studies, which have investigated imitation control, found engagement of mPFC and rTPJ, and they all had small sample sizes (between 10 and 20 participants). It is, therefore, unlikely that the inclusion of an additional study with a relatively small sample size (Brass et al., 2009) would change the results of the meta-analysis, given that they are weighted by sample size.

A further consideration is the relative size of the MD and ToM networks that we used in our analyses. Given that the MD network spans a much larger area than the ToM network, our analysis may be biased toward finding results in the MD network over the ToM network. Although this is true in a relative sense, we do not feel that it hinders our interpretation of the results in the ToM network for several reasons. First, regions of interest in the ToM network were not particularly small areas. The mPFC regions included several portions of the dorsal, middle, and ventral mPFC, and the rTPJ covered a relatively large area of cortex. Second, both networks were defined accurately based on prior work, which used large samples of participants. Thus, even though ToM areas were comparatively smaller than the MD network, they still covered a swath of cortex in regions functionally and precisely defined as the ToM network. Consequently, we feel confident that had these regions been consistently engaged across studies, we would have been able to detect them. Third, even if we only use CIMs across the whole brain, which report activation peaks from prior studies, thus avoiding issues with thresholding or choice of masks, we still do not find evidence for engagement near rTPJ and mPFC for the imitative compatibility effect (Figure 4).

An additional possibility to consider is that the difference between the results in terms of domain-specific and domain-general network engagement could be due to the differences in stimuli used in the studies included in the meta-analysis. However, the tasks are all conceptually, visually and cognitively similar to each other with only minor differences across all studies. For example, in Darda et al., (2018; Exp1 and Exp2), the stimuli consist of index and middle finger movements, whereas in Wang et al. (2011), hand opening and closing movements are used. Moreover, a recent meta-analysis also showed that behavioural performance is consistent across a range of studies that cover a range of minor methodological differences (Cracco et al., 2018). Given the lack of substantial differences between the studies and the consistent pattern of behavioural data, it seems unlikely that small differences could be responsible for these effects.

Finally, we acknowledge that fMRI is only one form of measurement, and it is important to consider how these findings mesh with results from other neuroscience techniques. For instance, neurostimulation studies have implicated rTPJ in imitation control (Santiesteban et al., 2015; Bardi et al., 2017). Using repetitive transcranial magnetic stimulation (TMS), dampening of activity in the rTPJ interfered with imitative, but not spatial responses (Hogeveen et al., 2014; Sowden & Catmur, 2015), whereas excitatory stimulation of the rTPJ by anodal transcranial direct current stimulation (tDCS) caused increased performance on the imitation task (Santiesteban et al., 2012). Further, in patients with lesions in the temporoparietal junction area, imitation inhibition deficits have been found to correlate with deficits in visual and cognitive perspective taking tasks, further supporting the role of rTPJ in imitation control (Spengler et al., 2010). Thus, there seems to be a discrepancy between neurostimulation and patient studies, and results from the current meta-analysis of fMRI studies. The evidence from neurostimulation and patient studies for the engagement of rTPJ in imitation control is, however, limited to a few studies with small sample sizes. Under any yardstick, therefore, the sum total of evidence from neurostimulation and patient studies can only be judged to be suggestive at present. It is based on a few studies with small sample sizes that lack formal power analyses and replications. Therefore, for more confirmatory evidence, future investigations with pre-registered and adequately powered replications are essential (Munafo et al., 2017; Zwaan et al., 2018; Nelson et al., 2018). In addition, it is also possible that the role of rTPJ in imitation control cannot be captured by univariate measurements and a more complex neural organisation is at play during imitation control.

### Theoretical implications

The lack of consistent activation in mPFC in the current meta-analysis and a difficulty in interpreting the role of rTPJ have implications for “specialist” theories of imitation. “Specialist” theories suggest that based on a dedicated neural circuit for social cognition, self-other control is crucial for the regulation of imitation, empathy, autism, and theory-of-mind (Brass et al., 2009; de Guzman et al., 2016; Sowden & Shah, 2014). However, more recent behavioural evidence suggests that imitation may not vary as a function of autistic-like traits or empathy, thus questioning the reliance of imitation inhibition on a distinctly social mechanism (Butler et al., 2015; Cracco et al., 2018; Genschow et al., 2017). Instead of a distinctly social mechanism, imitation control may involve domain-general cognitive control mechanisms, which are also engaged during the control of other non-social pre-potent response tendencies (Heyes, 2011; Cooper et al., 2012). Indeed, the dual-route model of automatic imitation proposed by Heyes (2011) can explain the control of automatic imitative tendencies without assuming a reliance on a self-other distinction. The model suggests that like other stimulus-response compatibility tasks, imitation control is mediated by long-term stimulus-response associations which are a product of learning. In line with this, the computational model put forth by Cooper et al. (2012) further substantiates this notion by demonstrating that spatial and imitative compatibility effects depend on similar cognitive processes, and any behavioural differences are accounted for by different sets of input nodes for spatial and imitative effects in a general dual-route framework (but see Berthental & Scheutz (2013) for a critique of this model).

Even though it is possible that imitation and spatial compatibility rely on a partly shared set of cognitive processes, this does not address the question of whether these processes also rely on similar or distinct neurobiological mechanisms. The current meta-analysis suggests that the selection mechanism in imitation inhibition is guided by a domain-general multiple demand system, which is also engaged during the inhibition of other non-social external influences. However, a lack of engagement of mPFC (and possibly rTPJ) in imitation control does not imply that they do not also play a regulatory role in imitation control. For example, mPFC has been demonstrated to exert a top-down influence during modulation of imitation via direct gaze (Wang et al., 2011). In addition, rTPJ showed a higher response when an interaction partner was believed to be human and looked human compared to when these animacy cues were absent (Klapper et al., 2014). These findings suggest that mPFC and rTPJ may play a regulatory role in imitation control and may be functionally connected to other networks without being directly engaged (Burnett & Blakemore, 2009). The current findings suggest that future work should postulate and test more complex models of imitation control, which extend beyond the operations of the theory of mind network.

In a similar manner, other socio-perceptual circuits, which extend beyond the MD network, may also be involved when inhibiting automatic imitative tendencies. In this regard, it is important to note the distinction between input- and mechanism-specificity. Of course, the input in the imitation inhibition task can be readily identified as emanating from a social entity i.e. a human hand. Thus, the observed input is clearly social in the sense that the observed agent offers opportunity for social interaction. Although the perceptual input is social, a domain-general selection mechanism may still operate in imitation control. Indeed, it is possible that the same selection mechanism operates across both social and non-social contexts. In the context of imitation, therefore, domain-specific action observation and person perception networks may functionally interact with domain-general control mechanisms in the MD network (see Figure 1). Thus, similar to other domains of social information processing, an interplay between domain-general and domain-specific networks may result in the control of automatic imitative tendencies (Baldauf & Desimone, 2014; Spunt & Adolphs, 2017; Zaki et al., 2010). Thus, the engagement of domain specific and domain general neural networks in imitation control may be more complicated that what current models of imitation suggest. Consequently, theories that move beyond a neat division and posit links between domain-general and domain-specific systems in imitation control need to be given greater emphasis in future work (Barrett, 2012; Spunt & Adolphs, 2017; Michael & D’Ausilio, 2015; Binney & Ramsey, 2019).

In conclusion, the current meta-analysis provides evidence that the selection mechanism when inhibiting automatic imitative tendencies is guided by the regions of the domain-general multiple demand network rather than a domain-specific system related to social cognition. Our meta-analysis questions the role of mPFC and right TPJ in imitation control, and suggests that current neurocognitive models of imitation control need further revision in order to account for the more complex nature of functional interplay between domain-general and domain-specific systems.

## Data and code availability statement

Database of co-ordinates and code used to perform the meta-analysis are available online at: https://osf.io/dbuwr/

## Ethics statement

The current work was a meta-analysis of existing data and there were no ethical considerations.

